# TULIP — a Transformer based Unsupervised Language model for Interacting Peptides and T-cell receptors that generalizes to unseen epitopes

**DOI:** 10.1101/2023.07.19.549669

**Authors:** Barthelemy Meynard-Piganeau, Christoph Feinauer, Martin Weigt, Aleksandra M. Walczak, Thierry Mora

**Affiliations:** Computational and Quantitative Biology, LCQB UMR 7238, Institut de Biologie Paris Seine, CNRS, Sorbonne Université, Paris 75005, France; Department of Computing Sciences, Bocconi University, Milan 20100, Italy; Laboratoire de Physique de l’Ecole Normale Supérieure, ENS, Université PSL, CNRS, Sorbonne Université, Université de Paris Cité, F-75005, Paris, France

## Abstract

The accurate prediction of binding between T-cell receptors (TCR) and their cognate epitopes is key to understanding the adaptive immune response and developing immunotherapies. Current methods face two significant limitations: the shortage of comprehensive high-quality data and the bias introduced by the selection of the negative training data commonly used in the supervised learning approaches. We propose a novel method, TULIP, that addresses both limitations by leveraging incomplete data and unsupervised learning and using the transformer architecture of language models. Our model is flexible and integrates all possible data sources, regardless of their quality or completeness. We demonstrate the existence of a bias introduced by the sampling procedure used in previous supervised approaches, emphasizing the need for an unsupervised approach. TULIP recognizes the specific TCRs binding an epitope, performing well on unseen epitopes. Our model outperforms state-of-the-art models and offers a promising direction for the development of more accurate TCR epitope recognition models.

## I. INTRODUCTION

T cells detect foreign invaders such as viruses, bacteria and cancer cells through their membrane-bound T-cell receptor (TCR), which recognize specific epitopes presented on the surface of infected or tumor cells. Epitopes are short (8-17 amino acid) peptide fragments presented by the major histocompatibility complex (MHC) on the surface of presenting cells, which are bound to by the TCR (Fig. 1A). The TCR is a heterodimer composed of the alpha and beta chains, which are coded by separate genes that randomly recombine during thymic development, giving rise to a large diversity of possible TCRs. Binding between the TCR and the peptide-MHC (pMHC) complex is highly specific [1, 2] and plays a key role in the activation of the adaptive immune response. Predicting pMHC-TCR binding from their amino-acid sequences is an important challenge in immunology. It has important applications to diagnostics, cancer immunotherapy, and vaccination, including the engineering of TCR against target antigens [3], or the design of optimized antigens in personalized cancer vaccines [4].

**FIG. 1.**
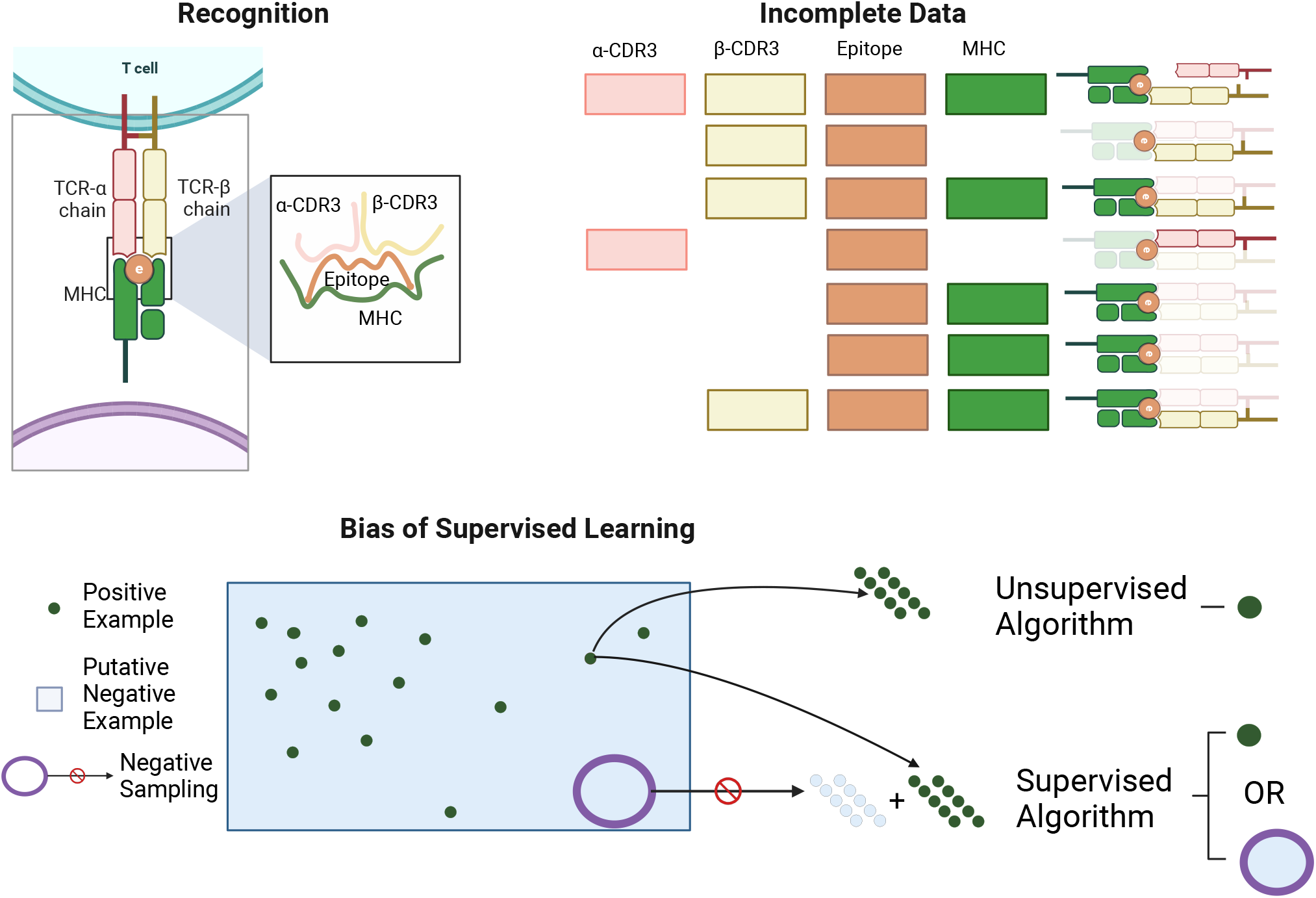
A Recognition: The TCR is composed of an *α* chain and a *β* chain, each one interacting with the epitope through its CDR3. The epitope is presented by the MHC. B Incomplete Data: Schematic representation of the current state of data availability for this binding problem. C The bias of Supervised Learning: Comparison of Supervised and Unsupervised approach. The unsupervised approach is only seeing positive pairs, it will only learn to recognize the specific signal of interacting pMHC-TCR. On the contrary, the supervised approach needs to sample negative examples, and the model will also try to capture the signature of none interacting pMHC-TCR. This may introduce a bias as the model can be learning to recognize some specific signal coming from the method used to generate the negative examples. Created with BioRender.com contained in incomplete datasets.

Given the difficulty to predict the structure and binding interface of pMHC-TCR pairs, predicting their binding affinity from general rules of protein interactions remains a promising but arduous approach [5, 6]. Recent experimental advances [7, 8] have allowed for the generation of an increasing amount of data linking TCR sequences to peptide-MHC (pMHC) complexes, providing a large number of binding pairs. These data are gathered in several freely available databases: VDJdb [9], IEDB [10], McPAS-TCR[11]. However, the number of possible 7-16 amino-acid peptides is very large, and the potential number of possible TCRs even larger (*>* 10^60^ [12]), meaning that experiments may only assay a small fraction of possible pairs. This calls for machine-learning methods capable of predicting the binding properties of unobserved pairs from a limited set of training data, by learning general rules of pMHC-TCR interactions.

Several studies have attempted to predict TCR specificity from sequence using a variety of machine learning techniques (see [13] for a recent benchmark), including deep convolutional networks (NetTCR2 [14]), decision trees and random forests (SETE [15], TCREX [16]) Gaussian process classification (TCRGP [17]), distance-based methods (TCRdist3 [18]), and language models (TITAN [19], Pan-Pep [20], ERGO2 [21], STAPLER [22]), and ensemble methods of Convolutional neural networks (DLpTCR [23]). Many approaches are inherently incapable of, or show poor performance at, predicting TCR affinity to epitopes that were not present in the training set (unseen epitopes), either by design or by lack of generalizability across epitopes [19, 22]. This fundamentally limits their applicability, in particular in the context of cancer neoantigens which are often unique to each patient.

Existing models are often trained on a subset of all available data, because of requirements on quality and consistency. Experiments rarely report all four elements of the binding complex: the peptide, the MHC, the alpha and beta chains of the TCR (Fig. 1B). Because information about pMHC specificity is shared across both chains [7, 24], many methods choose to focus on data that report both chains, leaving out the large amount of information Another limitation of existing approaches is that they treat the binding prediction as a supervised learning task, which requires both positive and negative examples to train a binary classifier. However, the biological data at our disposal is not of this type, consisting only of positive examples. To address this issue, negative examples are often generated using random association, but these can lead to subtle biases [22]. The fraction of random pMHC-TCR functional associations is estimated to be *≈* 10^*−*6^–10^*−*4^ [25], meaning that non-binding pairs widely outnumber binding ones. Therefore sampling the negative space properly for training a supervised classifier is difficult. Using a supervised approach may push models to learn the biases in the negative data provided, rather than biologically meaningful patterns.

The case of having only data from one class is usually called One Class Classification (OCC), and is not new in biology [26]. Generative models are one solution to tackle this task, as we do not need any negative example to train it [27].

In this paper we present Transformer-based unsupervised language modeling for Interacting pMHC-TCR (TULIP-TCR), an encoder-decoder language model, which addresses these limitations. The model is flexible, leveraging all possible data sources regardless of their quality or completeness and including single-chain data, but also learning useful representations of the TCR and epitope space from examples where only one of them is present. The approach is unsupervised in the sense that we do not predict explicitly a binary variable that indicates binding, but a probability score trained only on interacting sequence pairs. This allows us to avoid the pitfalls associated with creating artificial samples of non-interacting sequence pairs [27]. TULIP outperforms state-of-the-art methods on the most studied peptides for which data is abundant, and shows significant predictive power on unseen epitopes.

## II. RESULTS

### A. Model overview: a flexible and unsupervised architecture

Our model is inspired by techniques used in Natural Language Processing (NLP), where models are commonly trained on large text corpora [28]. In our approach, we adapt these techniques by replacing words with amino acids. The central concept behind our model is translation, which involves predicting the next token (word or word pieces in NLP, or amino acid in protein sequences) based on the previously generated tokens, as well as the source (sentence in NLP, or amino-acid sequence in proteins). During training, the model learns the patterns and dependencies that govern the relationships between tokens. By training only on positive examples, the model learns the rules that govern token ordering.

Models that predict each amino acid conditioned on the previous ones are called autoregressive. This allows us to compute the probability of each sequence As 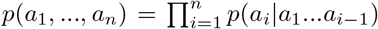, and to efficiently sample new sequences, with tremendous recent success in modeling language [29] .

The training process involves maximizing the conditional likelihood of the observed sequences (positive pairs), effectively defining a probability distribution over the space of sequences. As a result, the model is trained to assign higher probabilities to positive pairs (binding pairs) compared to negative pairs (non-binding pairs) without having been trained on any negative pairs.

Our model uses the Transformer architecture, specifically the encoder-decoder variant originally developed for translation tasks [30]. In this architecture, the encoder receives a protein sequence as input (a sentence in the source language in NLP), and the decoder aims to generate an interacting protein sequence (the translated sentence in NLP) as its objective. The decoder coupled with the encoder is an autoregressive generative model, which defines the conditional probability distribution of the output given the input. The encoder-decoder approach has been successfully applied to investigate interacting amino acid sequences [31, 32].

Our problem implies interactions between 4 elements: the epitope, the MHC, and the alpha and beta chains of the TCR. We reduce the chains to their third complementarity determining regions (CDR3) known to be primarily contacting the epitope [33]. We denote the *α*-CDR3, *β*-CDR3 and epitope sequences as 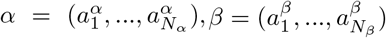 and 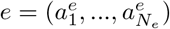. We extend the existing architecture and define 3 encoders and 3 decoders for the *α*-CDR3, *β*-CDR3, epitope sequences and a special embedding layer for the MHC, which we treat as a categorical variable MHC (its protein sequence is ignored, as we expect only the MHC class to be relevant). The details of this architecture are shown in Fig. 2A. Each model takes the MHC and the 3 chains as input, and try to predict each chain given the two other ones and the MHC.

**FIG. 2.**
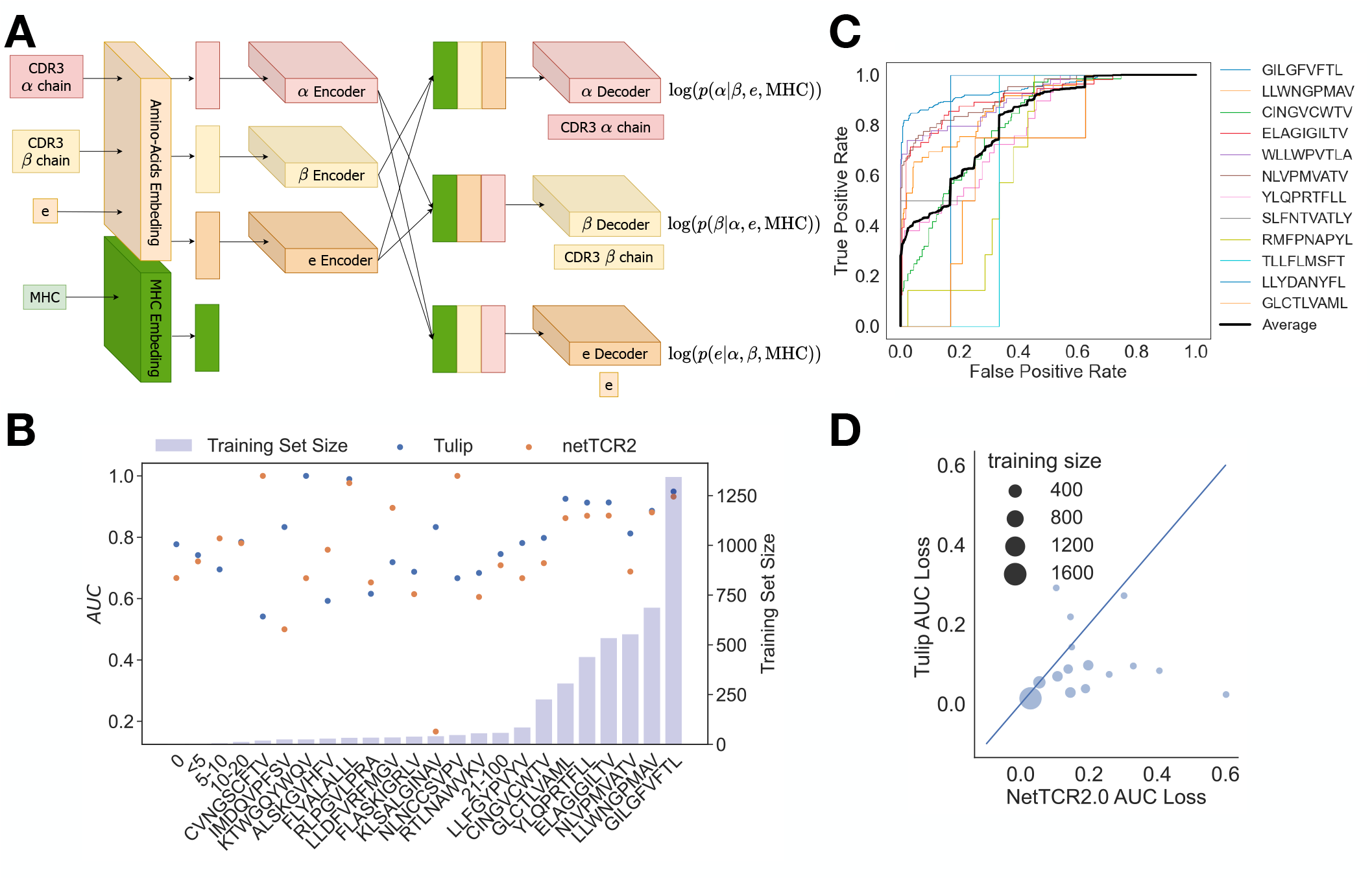
A - TULIP architecture: Amino acids of each chain are embedded, then encoded by its chain-specific encoder. The MHC is also embedded. The MHC embedding and the encoded chains are then concatenated (all except the embedding of the sequence to decode) and given to the decoders. The decoders are then modeling the conditional probabilities of each chain given the MHC and the other available chains. B - Results of a finetuned TULIP on the most abundand peptides. Comparison is made with NetTCR2.0. C - ROC curve and PPV curves for the most studied peptide. The average of these ROC curves appears in red. D - We compare two ways of selecting the negative example. We compare the loss of performance of NetTCR2 and TULIP between an easy and a hard case of negative sampling. In the easy case the TCRs are randomly selected from the test set, whereas in the harder case, the TCRs are reweighted in order to have a uniform distribution over the true cognate epitope. This second choice removes the bias of having most negative examples using TCRs from the few highly over-represented peptides. Because TULIP is unsupervised it is more robust to change in the negative sampling.

They define conditional probabilities such as *p*(*e*|*α, β*, MHC). These conditional probabilities can be used to match interacting protein sequences [31], since pairs that bind are expected to have higher probabilities than non binding ones.

This model can be used with incomplete data by determining, e.g., restricted conditional like *p*(*e*|*α*) when the beta chain and the MHC class are not available. This flexibility enables us to use every known data source available for model training and for prediction. More details about the architecture and the training can be found in the Methods section IV B.

### B. Predicting new TCRs binding to known epitopes

We first evaluate the performance of TULIP for epitopes presented on the common HLA-A*02:01 allele, which is commonly used to assess such models [14]. We compare TULIP with NetTCR-2.0, a state-of-the-art supervised model [14]. We collected data for which the epitope, alpha chain, and beta chain were all present. We then created a random split of 85% for training and 15% for testing, excluding any sequences from training in which the TCR was also present in the test set. Negative examples were generated within each split by randomly pairing TCRs to a different epitope, and this process was repeated five times for each sequence. To avoid overlap between the training and test sets, negative examples were sampled within each split. We refer to this database, comprising both the training and test sets, as the Specialized Dataset (SD). NetTCR was trained on the training SD and its performance was evaluated on the testing SD for each epitope separately using the area under the curve (AUC) of the receiver operating characteristic (ROC) curve as a performance metric.

The primary aim of TULIP is to be trainable on a larger database. To ensure a fair comparison, we trained it using the following protocol: Firstly, we removed all sequences from the full database that shared the same alpha or beta chain as the test set of the SD. We trained the TULIP on this filtered dataset for 100 epochs and then fine-tuned it for an additional 40 epochs using the positive examples of the training SD. To compute the AUC, we approximated the probability of binding as log(*p*(*e*|*α, β*, MHC)) *−* log(*p*(*e*|MHC)) (see IV C), which quantifies the increase in the odds of observing *e* upon being recognized by the TCR.

We compared the performance of TULIP with NetTCR-2.0 on the testing set of SD, and computed the AUC separately for each epitope in Fig. 2B. Rare epitopes were grouped by similar training set size, and their AUC averaged. The results indicate that TULIP outperforms netTCR2.0 on almost all epitopes. For completeness, we plotted the ROC curves of TULIP in Fig. 2C. These curves reveal a very good performance on the top-ranked prediction as ROC curves start with a vertical line up to 0.5 of True Positive Rate before observing the first False Positives. This steep start is extremely interesting as it implies that the model is extremely good for the sample for which it is the most confident.

Because the AUC treats positive and negative examples symmetrically, it is particularly sensitive to the choice of negative samples, which the supervised method can exploit to artificially boost its performance [22]. To illustrate this bias, we implemented a different sampling approach for negative examples within our specialized datasets. Instead of uniformly sampling non-binding TCRs, we uniformly sampled another epitope and then selected one of its associated TCRs. This alternative sampling procedure aims to counteract the bias introduced by the over-representation of TCRs from the most commonly observed epitope in the negative sets, which leads supervised methods to learn the features of TCRs binding to that epitope, instead of learning the features of the positive TCRs. Fig. 2D shows that performances of both TULIP and NetTCR-2.0 decrease when this alternative sampling is applied, demonstrating the importance of this bias. This alternative sampling does not affect the TULIP model itself, whose training does not involve negative examples, but it does affect its AUC which relies on negative examples. However, the bias is more pronounced for a supervised method such as NetTCR-2.0, as evidenced by the fact that most points fall below the diagonal.

### C. Generalization to unknown epitopes

A major challenge of pMHC-TCR binding models is to be able to generalize, i.e. to make binding predictions on epitopes that were not used in the training set (unseen epitopes). This ability varies a lot depending on the considered epitope, notably as a function of how similar it is to other epitopes used during training, making comparisons between methods and different contexts difficult. Here, we propose a systematic approach for assessing generalization across thousands of unseen epitopes, by stratifying them according to their distance to the training set.

We split the full database into a test set comprised of epitopes with fewer than 20 examples, and a training set composed of those with more than 20 examples. TULIP was subsequently trained on the training set for 100 epochs, following which its performance was evaluated on the testing set, yielding 1796 AUCs. The Levenshtein distance between each unseen epitope and its closest counterpart in the training set was then computed. To mitigate potential bias from deep mutational scanning (DMS) experiments, which contain large numbers of closely related sequences, we identified TCRs that were associated to similar peptides and deleted them from the training set. For each peptide in the test set, the subset of peptides with minimal distance in the training set was identified, and all TCRs associated to them were removed from the test set. All TCR sequences associated with both the peptide from the test set and any peptide in the subset of closest peptides within the training set were removed from the training set. Despite these corrections, the dataset is still very biased. The distribution of TCR per epitope is skewed with a heavy tail (SI Fig. S1), and epitope representation is mostly biased towards COVID peptides and neoantigens (SI Fig. S2).

We computed the average AUC of epitopes as a function of their distance to the training set (Fig. 3A and SI Fig. S3, and SI Fig. S4 as a function of normalized distance). TCR similarity between the training and the testing sets is shown in SI Fig. S5. Machine learning methods tend to perform better in the region closer to its training set. It is a common phenomenon in all machine learning approaches for the model’s capacity to extrapolate and generalize to decrease as one moves further away from the training set. We used 3 different methods for sampling the negative examples in the model evaluation (Fig. 3B). In the Unseen Unconnected Random Association (UURA) and Unseen Connected Random Association (UCRA) methods, negative pairs are drawn by picking a random TCR and a random epitope that were not in the training set. In the UURA, which is more rigorous, the true cognate epitope of the picked TCR is also unseen, while in the UCRA it can be any epitope (seen or unseen). In the Healthy Repertoire Sampling (HRS), the TCR is chosen at random from the repertoire of healthy individuals (for which the epitopes are unknown) taken from Ref. [34]. The results obtained with the most conservative negative sampling procedure (UURA, in blue) indicate that TULIP shows good generalization for epitopes that are close to the training set. This performance decays quickly with distance, reaching 1/2 (chance level) around at an edit distance of around 4.

**FIG. 3.**
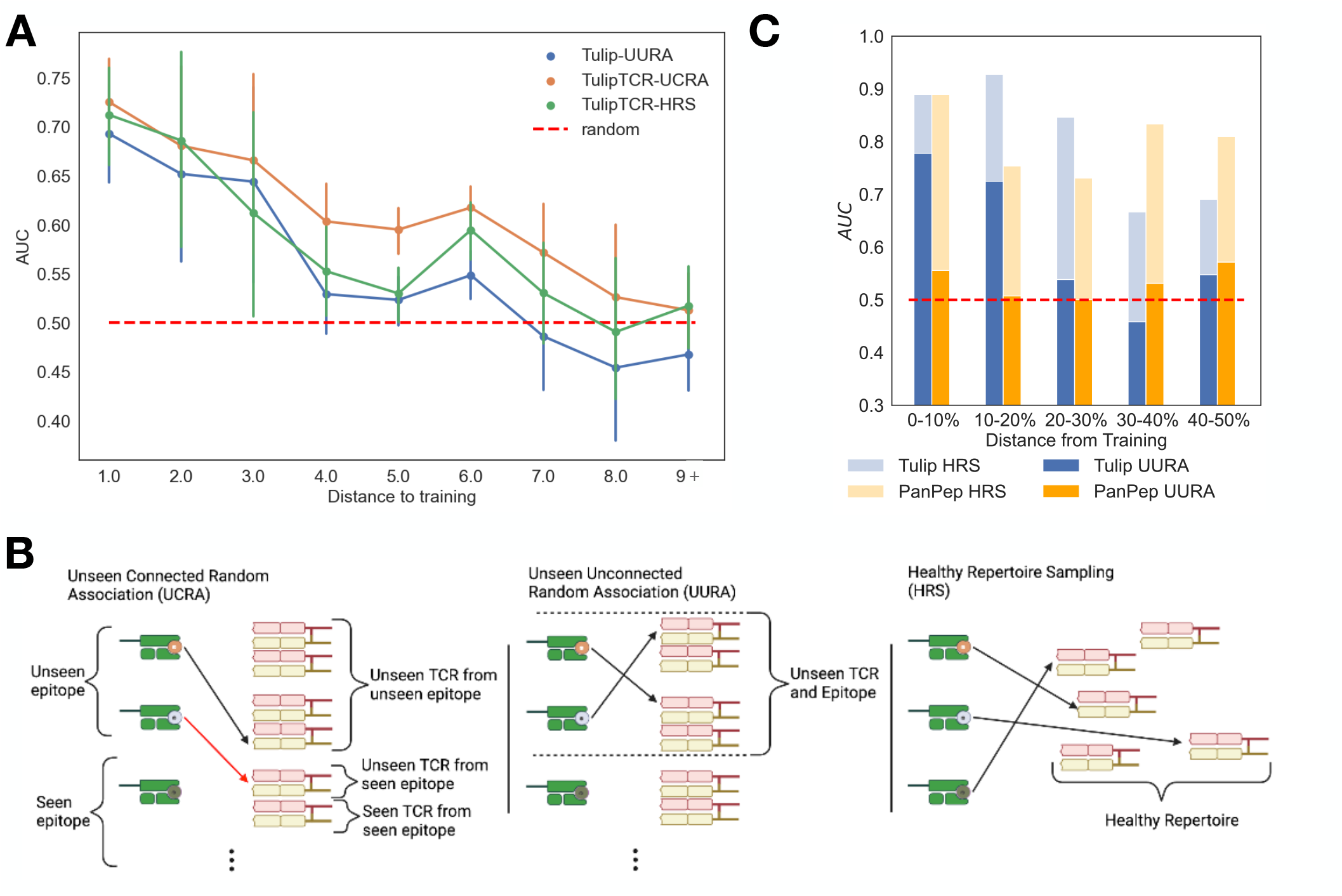
A. Performance of TULIP on unseen peptides as a function of the distance to seen peptides. Up to edit distance 4, a clear signal can be seen. This analysis is done on a large set of peptides (171 at distance 1, 43 at distance 2, 44 at distance 3, 161 at distance 4, 501 at distance 5, 500 at distance 6, 103 at distance 7, 54 at distance 8, 219 at distance 9 and more). We also illustrate the role of negative sampling by showing the performance with three different Negative sampling methods. The details of these methods are explained in B. Our unsupervised methods show less variability with respect to the sampling methods compared with other supervised methods as shown in C and SI Fig.S6 B - We detail here three different methods to sample the negative of unseen epitopes. We illustrate the fact that in the original data, several TCR can be binding a single epitope, by putting two TCR in front of each epitope in the plots. Unseen Unconnected Random Association: the epitope and the TCRs are unseen and the TCRs used for the negative are binding with an unseen epitope. Unseen Connected Random Association: the epitope and the TCRs are unseen and the TCRs used for the negative can be binding to any epitope. We emphasize in red the association with a unseen connectd TCR, as it is the difference with UURA. Healthy Repertoire Sampling: Negatives are sampled from a healthy repertoire. Created with BioRender.com C - Testing the effect of change of the negative sampling on unseen peptides for PanPep and TULIP. We realize that PanPep performance does not resist changing the negative sampling process for unseen peptides contrary to TULIP. For the HRS, we reused the negative example from the original paper.

For comparison, we also investigated the performance of existing models, PanPep [20], Ergo2 [21], and DLpTCR [23], but re-evaluated using the more rigorous UURA negative sampling method not used in the original studies (as the training/testing split of STAPLER [22] was not available at the moment of writing, we could not compare performance with that method). For instance, the performance of PanPep on unseen epitopes, which was originally assessed using the HRS method, drops to chance level when using the more stringent UURA (Fig. 3C). By contrast, TULIP, when tested on the same dataset (and re-trained on data that excluded that test set) retains some predictability. Similar results for DLpTCR are reported in SI Fig. S6A. We also compared our findings with ERGO2, which was trained using the UCRA method for negative sampling. Conducting a test by resampling the TCRs with the more conservative UURA shows that the resulting AUC also decays to values close to chance level (SI Fig. S6B). Note that STAPLER [22] was also evaluated using UCRA, potentially inflating its performance on unseen epitopes.

These findings underscore the risks of using negative samples during training. Since most pairs are negative, identifying a non-binding pair carries very little information. Any signal captured from negative examples is likely a result of batch effects introduced by the negative sampling procedure. This justifies the choice of an unsupervised architecture for pMHC-TCR binding. Thanks to this structure, TULIP is robust in the face of changes in negative sampling approaches, since training does not use any negative samples.

### D. Predicting the effect of neoantigen mutations on TCR activation

To further test our model’s ability to predict binding to different epitopes, and to predict epitope mutations that may evade immune recognition, we applied it to deep mutational scans of epitopes against fixed TCRs from [35]. A deep mutational scan of the epitope binding with a fixed TCR is a systematic analysis that explores the effects of multiple genetic mutations within the epitope on its interaction with a specific T-cell receptor. The study involved 6 deep mutational scans of two epitopes (HLA-A*02:01 restricted NLVPMVATV and IMDQVPFSV also present in the training set) against three TCR targets each. For each of the 19 *×* 9 single-amino acid variant of the epitopes, the affinity to the TCR was assessed by measuring the epitope concentration at which 50% of T-cells were activated in culture (EC50). Observing binding in an experiment requires both the binding of the peptide with the MHC, and of the TCR with the pMHC. We used the joint probability of binding as a score log *p*(binding(e *−* MHC), binding(e *−* TCR)|*α, β*, e, MHC) We approximate this quantity following the method in IV C by log *p*(*e*|*α, β*) *−* log *p*(*e*) + log *p*(*e*|*mhc*) *−* log *p*(*e*) as a predictor of this affinity. The comparisons between model and experiments are shown in Fig. 4A. Despite high variability, our model was able to capture the fundamental properties of binding in epitope space. To quantify performance, we measured the Spearman correlation between our score and the measured EC50. The score correlates up to 0.47 for the best TCRs. While predictability is limited, these results are encouraging considering that the model was trained on data with a large excess of TCRs relative to epitopes, and applied to data with a large excess of epitopes relative to TCRs. To assess how much of this predictability is due to peptide-MHC only (irrespective of the TCR), we compared these results with NetMHCpan [36], which is based solely on the epitope-MHC interaction, and found a lower correlation (SI Fig. S7). This highlights the importance of the TCR-epitope interaction in the experiment.

**FIG. 4.**
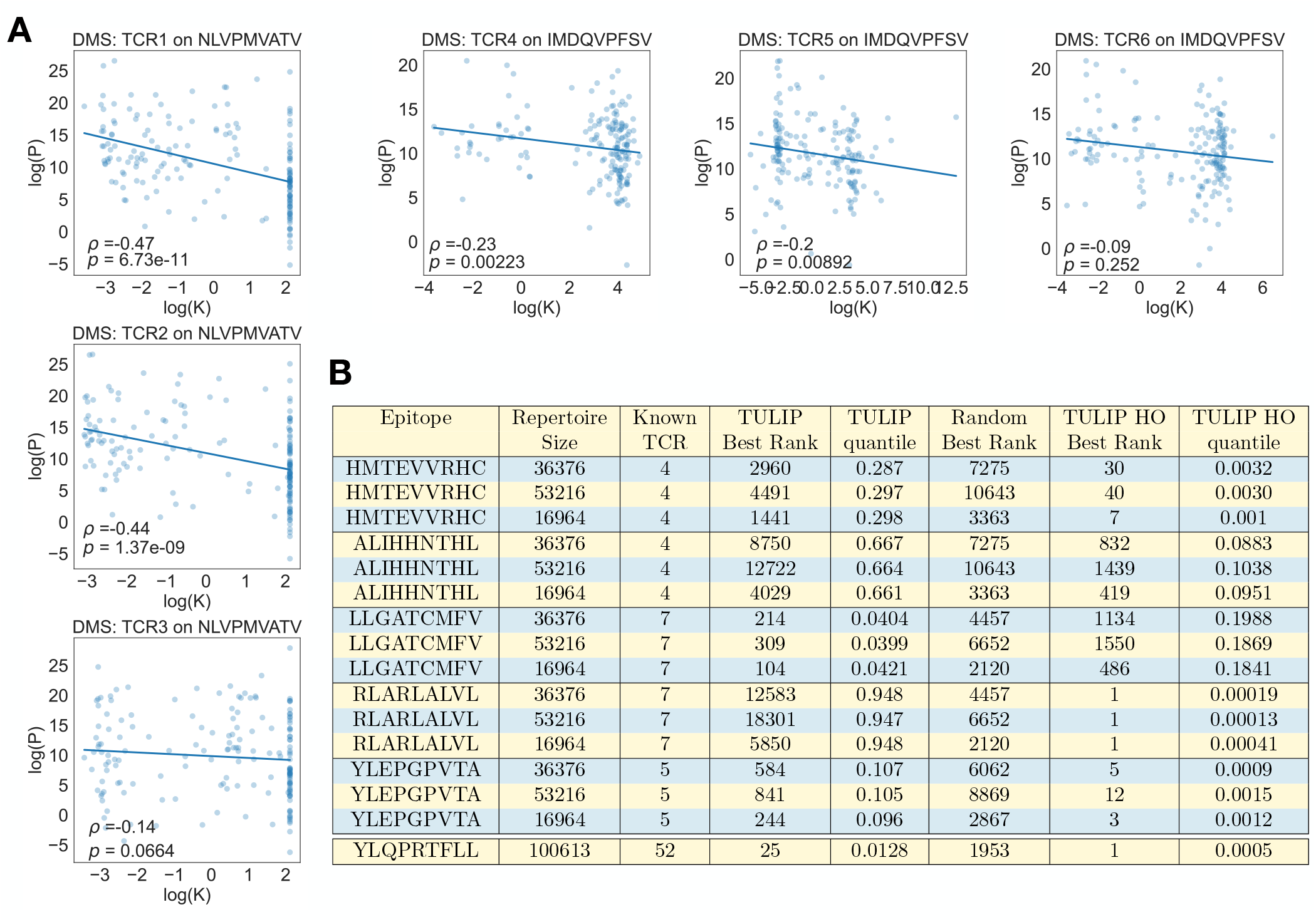
A. Effect of single epitope mutations on the TULIP score (log *P*) predicts TCR binding (dissociation constant *K* in *μ*g.ml^*−*1^) measured by deep mutational scan experiments [35]. The reported *ρ* and p-values correspond to Spearman correlations. B. Repertoire mining for neoantigen-binding TCRs. The TCR repertoires of 3 healthy HLA*A02:01 donors from [34] were spiked with TCRs known from the literature to bind to 5 neoantigens. Sequences from the augmented repertoires were ranked by the model according to their predicted affinity to the neoantigen of interest. Reported is the rank of the best-scoring neoantigen-binding TCR. The p value corresponds to the probability of achieving that rank by chance. Two training procedures were used: one where all TCRs associated to the neoantigen of interest and related peptides were removed from the training set (TULIP), and one where only the TCR to be ranked was removed (leave-one-out; TULIP LOO). We also integrated the quantile of our prediction under the null model. This is easily interpretable as the probability that a random model would achieve equal or better performance than TULIP.

### E. Repertoire mining for neoantigen recognition

We then asked whether the model could pick TCRs binding to a particular epitope from whole repertoires. We focused on TCRs binding to 6 HLA-A*02:01 restricted epitopes, including 5 cancer-associated neoantigens (Cyclin D1: LLGATCMFV [37]; p53: HMTEVVRHC [38, 39]; HER2: ALIHHNTHL [40]; TPBG: RLARLALVL[41]; and gp100: YLEPGPVTA [42], see Table S1 for the full list). In addition, we looked for TCRs specific to the SARS-CoV-2 spike protein epitope YLQPRTFLL in the CD8^+^ repertoire of a COVID-19 infected donor at the peak of the response at day 15 [43]. YLQPRTFLL-specific TCR harbored by the same donor were identified in a separate study using a multimer-binding assay [44].

We first considered a scenario where no prior knowledge about TCRs binding the epitope was available, by removing these entries from the training set, as well as all TCR-epitope pairs whose epitope is similar to the epitope of interest (less than 4 amino acid substitutions). For the SARS-CoV-2 epitope, we also removed all TCRs associated with YLQPRTFLL as well as similar epitopes (‘YLRPRTFLL’ and ‘YYVGYLQPRTFLL’) to mitigate potential leakage effects. In the second scenario (hold out, or HO), we only removed from the training set the TCR that we want to find: one neoantigen-associated TCR at a time in the case of neoantigen, and all epitope-specific TCR from the donor as reported by the multimer assay in the SARS-CoV-2 repertoire. In both cases, we removed redundancies of the alpha and beta chains: when several TCRs has the same alpha chain, we only retained one of them, and likewise for beta chains.

For each neoantigen, we mixed neoantigen-associated TCRs (all of them in the first scenario, and only the removed one in the LOO scenario) with 3 unrelated TCR*αβ* repertoires of HLA*A02:01 positive donors from Ref. [34]. For the SARS-CoV-2 epitope, we simply considered the CD8^+^ TCR*β* repertoire at day 15 from [43]. We then asked TULIP to rank each TCR according to the predicted binding to the epitope of interest. The results, reported in Fig. 4B, show that TULIP in many cases narrows down the list of candidate TCR to a relatively small number, even when it was trained with no knowledge about the neoantigen-associated TCRs. When it does (TULIP LOO first index column), it can even identify the neoantigen-associated TCR within the very best ranked ones. In the case of the SARS-CoV-2 epitope, performance was excellent even when in the first scenario (no prior knowledge about the epitope), and perfect (best rank 1) in the hold-out scenario.

That analysis focuses on the top-ranking TCR for each neoantigen, emphasizing precision in detecting potent binders within the repertoire. This deliberate emphasis on the upper tiers of the score distribution provides insights into the model’s discriminative power and its ability to identify TCRs with high binding affinity to specific epitopes.

## III. DISCUSSION

In this study, we have presented a novel approach for TCR-epitope binding prediction that overcomes key limitations of current methods. We demonstrated the model’s ability to generalize to unseen epitopes, which is a critical factor in real-world applications where the specific epitope of interest may not be known in advance. Furthermore, we addressed the recurrent bias that can arise from using negative examples generated through random pairing in previous supervised approaches. To mitigate this bias, we proposed an unsupervised learning framework that trains the model exclusively on positive examples, allowing it to focus on recognizing patterns within these interactions.

The elimination of negative examples in our approach was driven by the recognition that randomly generated negative examples can introduce biases, potentially compromising the model’s predictive accuracy. By training solely on positive examples, our model avoids such biases and can more effectively capture the specific signal of interacting pMHC-TCR complexes.

One difficulty in evaluating and comparing methods is that the exact TCR-epitope binding prediction task may differ across studies and applications. For instance, looking for epitope-specific TCR within the peripheral repertoire is a different task than finding them within responding clones in lymph nodes or in tumor tissues. Likewise, identifying TCRs binding to a neoantigen but not to the wildtype is not the same as identifying the response to a specific antigen within a repertoire. Some of the biases discussed earlier arise from unclear or unrealistic definitions of the tasks. When the objective is to recognize patterns in binding complexes, the unsupervised approach emerges as the more natural choice. Supervised approaches can only demonstrate their potential in specific use cases where negative samples can be precisely defined (e.g. sorting cells that do not carry an activation marker or do not bind a tetramer, although these negative examples are typically not reported in studies). Careful consideration should also be given to the sampling of negative examples. Negative examples should be selected to be close enough to the classification boundary, making them challenging examples (referred to as Hard Negative Sampling). The combination of these constraints, including a well-defined and restricted negative subspace, the difficulty of examples, presents significant challenges for most use cases, lead us to conclude that unsupervised approaches should be preferred for most applications.

We emphasize the importance of utilizing all available data sources, regardless of their completeness or quality. The same is true for NLP approaches, which usually start by collecting and training on as much data as possible. The TCR-epitope binding prediction task often suffers from the scarcity of comprehensive data, as obtaining complete TCR sequences along with corresponding epitopes and MHC information is challenging. However, our approach is designed to be flexible, leveraging the available data and accommodating situations where only partial data is accessible. By using both alpha and beta chains when available, while being able to learn from one chain alone, our model can make the most of the data at hand and extract valuable insights.

While our proposed model shows promise, it is essential to conduct fair and rigorous model comparisons to assess its performance accurately. The field of TCR-epitope binding prediction often lacks standardized benchmark datasets and evaluation protocols (but see [13]), leading to difficulties in comparing different models. To address this challenge, future research should focus on establishing standardized benchmarks and evaluation procedures that encompass diverse datasets and evaluation metrics beyond classification.

One limitation of our approach is that the model yields only probabilities of pairs of sequences, rather than a proper binding constant, for which titration data (where the concentration of the epitope is varied) would be needed. Another limitation is that large areas of the epitope space have not been measured, and some parts are extremely hard to measure. For example, having a model able to determine the risk that a TCR binds to self-proteins, would be extremely useful for predicting the safety of T-cell therapy, but such TCRs are by construction hard to observe, and the lack of data is a major limitation for further progress in this direction.

Since our model is generative in nature, it would be interesting to experimentally test its ability to generate de novo TCR sequences for given epitopes, or for combinations of related epitopes to which it would be cross-reactive. This avenue of research could provide valuable insights into the design and discovery of TCRs with specific binding capabilities.

## IV. METHODS

### A. Data collection

Data acquisition in the field of immunology presents a major challenge. The intricate process of T cell receptor (TCR) binding to its respective epitope depends on four critical elements: the epitope itself, the major histocompatibility complex (MHC), and the alpha and beta chains of the TCR. While each component has been extensively studied in isolation, the number of instances where all four components are jointly available remains remarkably scarce.

In this study, we present a novel computational aiming approach at constructing a model capable of learning from incomplete data. To achieve this goal, we curated data from multiple sources, maximizing the total sample size at our disposal. Specifically, we first accessed the VDJdb database [9] in its entirety, which boasts the highest data quality among our available resources (see Table I).

**TABLE I.** Summary of the data sources used for training.

|  | VDJdb<br>epitope<br>with MHC | VDJdb<br>epitope<br>without MHC | McPAS-TCR<br>epitope<br>with MHC | McPAS-TCR<br>epitope<br>without MHC | IEDB<br>epitope<br>with MHC | IEDB<br>epitope<br>without MHC | netMHC<br>epitope<br>with MHC |
| --- | --- | --- | --- | --- | --- | --- | --- |
| Alpha and Beta | 29251 | 0 | 5021 | 77 | 5021 | 77 | 0 |
| Alpha alone | 6750 | 0 | 1065 | 87 | 1065 | 87 | 0 |
| Beta alone | 23011 | 0 | 9898 | 218 | 145332 | 11 | 0 |
| No TCR | 0 | 0 | 0 | 0 | 451 | 11 | 663767 |

We also added the IEDB database of Tcell receptors and McPAS-TCR dataset [10, 11] (see Table I).

The IEDB database is more diverse but we observed a much poorer quality of the data, and there was never used for finetuning.

This accounts for 209779 not redundant data points containing the epitope and at least one chain of the TCR.

The instances listed above consist of T-cell receptors along with their respective epitopes and major histocompatibility complex (MHC). Regrettably, the MHC or one of its two chains is frequently absent. Additionally, the diversity of epitopes is relatively low compared to that of TCRs, with each epitope possessing multiple T-cell receptors.

To supplement our data, we incorporated the training database of netMHC, which is solely composed of MHC and epitope information. Although this dataset does not directly aid in comprehending the correlation between TCR and epitope, it is advantageous in two ways. Firstly, the dataset encompasses a wide range of epitopes, which assists the model in comprehending the true diversity of potential epitopes. Secondly, in order to achieve effective transfer learning between MHC, the model must comprehend what is distinct to each MHC and what can be transferred. Therefore, the netMHC database aids in better modeling the specific role of MHC in the epitope modeling process. We gather 663, 767 peptides with their MHC (see Table I).

We gathered all this data in a single one that we will refer to as the Full Dataset (FD).

### B. Model definition

Our model is an extension of the well-known Transformer model, in its encoder-decoder version. In the original version [30], the method was used for translation. During training the encoder was given a sentence in the source language and the decoder was given the translation in the target language as an objective to produce.

In our specific problem, we would like to condition our model on more than one interacting element. We, therefore, need to extend the existing architecture. We define 3 encoders, 3 decoders, and two embedding layers: an *α*-encoder that is specialized in encoding the *α*-CDR3, an *α*-decoder that is specialized in decoding the *α*-CDR3, a *β*-encoder that is specialized in encoding the *β*-CDR3, a *β*-decoder that is specialized in decoding the *β*-CDR3, an epitope-encoder that is specialized in encoding the epitope, an e-decoder that is specialized in decoding the epitope and finally an amino acid embedding and an MHC embedding, (as we decided to represent the MHCs as categorical variables). First experiments on initializing the decoders with the weights of pretrained general purpose proteins masked language models did not show any sign of improvement. TCRs *α* and *β* chains exhibit unique characteristics and patterns that are distinct from general protein sequences. The core of the loops of the CDR3 is extremely variable. On the other hand, epitopes are much smaller than usual proteins and presented inside an (MHC). All these factors imply that general rules for proteins do not transpose easily to our scenario. By utilizing dedicated encoders and decoders tailored to the specific nature of TCRs and epitopes, we can capture and encode their domain-specific features more effectively. This specificity enables the model to focus on relevant information and potential interactions specific to TCR-epitope binding.

While we refer to the original work on Transformer [30] for precise details on the attention layers and the encoder-decoder architecture, we review here the key components. Sequences are encoded by their specific encoder and used as the input for the decoders. They are processed through alternating blocks of self-attention and linear layers.

Typical vocabulary sizes in NLP are in the order of 10^4^ to 10^5^, while in our case we have a vocabulary *V* is composed of the 20 amino acids and some special token (PAD for padding, EOS for End-od-Sentence, SOS for Start-Of-Sentence, UNK for the Unknown characters). The sequence embedding is composed of two parts, one for the amino acid identity and one for the position in the sequence. We learn a dictionary, mapping each of the amino-acids to a vector of dimension *d*_*model*_. The sequence position is embedded as a vector in the same way, learning an embedding vector for every position. We also learn a specific embedding for the most common HLA types. The embedding of each sequence is taken as the sum of the amino-acids and positional embeddings. The embedded amino-acid sequences are then passed to the respective encoders, mapping them to a latent representation *z*^*T*^ = (*z*_1_, …, *z*_*n*_), *T ∈* (*α, β, e*). The encoded sequences are then concatenated with the MHC embeddings before being sent to the decoder. For each decoder, we concatenate the MHC embedding with all the encoded sequences except the one that the decoder will reproduce, as we do not want to give a decoder the sequence it is supposed to reproduce. The details of these groups can be seen in Fig2. For example, we concatenate the encoding of the *α*-CDR3, the *β*-CDR3, and the MHC for the Epitope decoder, as the epitope decoder should be conditioned on everything but himself. The decoders are then trained to predict their respective amino-acid sequences conditioned on the encoded pieces of information.

The decoder implements an auto-regressive distribution, for example

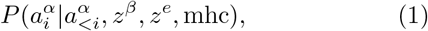

defining the probability of the *i*^*th*^ amino acid in the *α − CDR*3 sequence given the preceding amino acids 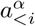 and the hidden representation of the others elements. During training, we use the true amino acids for 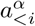. This way of predicting the next amino acid in a sequence is called Causal Language Modeling (CLM). The loss associated with this task for a single sequence is simply the cross entropy for every predicted token.

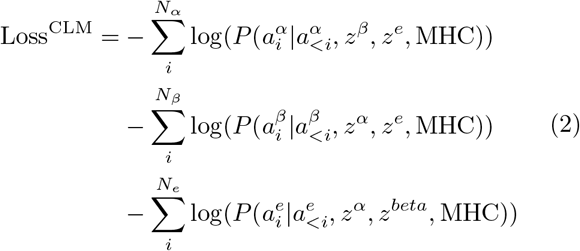

We schematize the forward pass of Tulip in the following pseudo-code:

#### Algorithm 1

TULIP

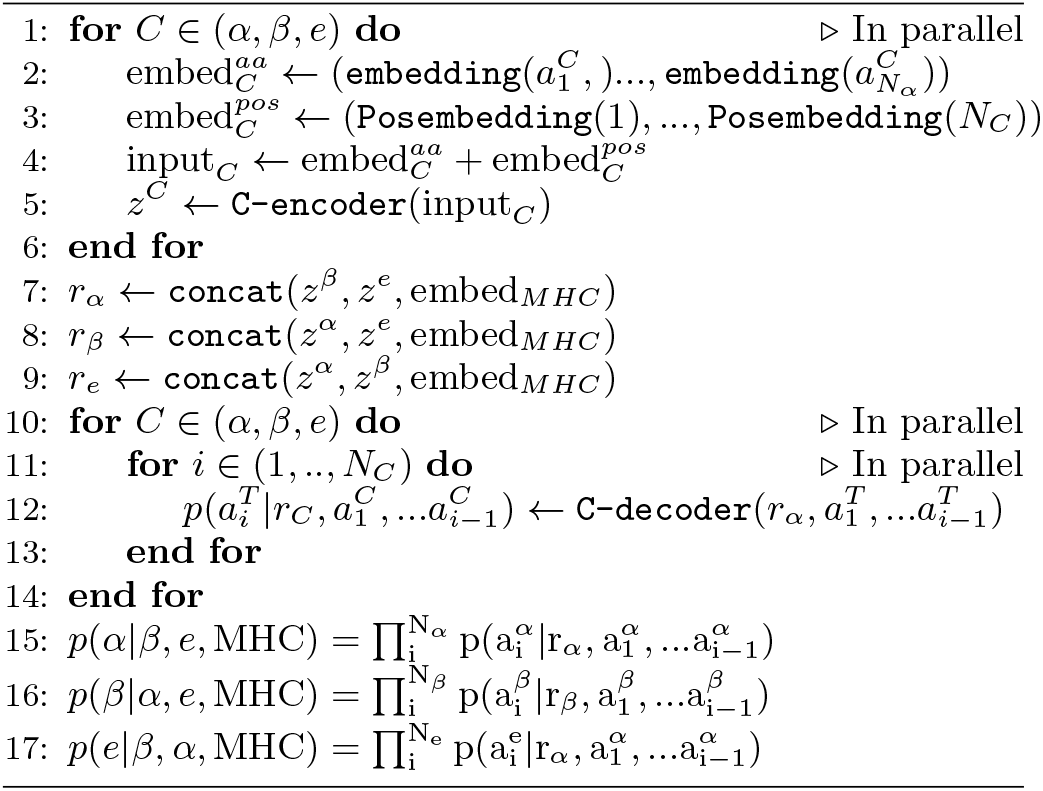

This approach has already been used for proteins in many works. Especially in [31] an encoder-decoder model was used to investigate interacting amino-acid sequences. The first thing to remark is that the decoder defines autoregressively a probability distribution over the generated sequence. It is generative as we can sample new examples but if we give it an existing specific sequence it will give us its probability. When coupling this to an encoder the probability distribution becomes a conditional probability distribution (conditioned on the input of the encoder). These conditional probabilities can be used for matching interacting protein sequences [31].

One interesting property of the Attention mechanism of the transformer is that it is position-blind and flexible with respect to the length of its input. This implies that it does not hard code in its weights where it is expecting to find specific elements. If a chain is missing, let’s say the *α*-CDR3, we can only gather the MHC and the *β*-CDR3 before giving it to the epitope decoder. The encoded *β* amino acids end up in the first position of the gathered encoding. This is not a problem thanks to the position-blindness of the encoder-decoder attention. To be more precise, the missing *α*-CDR3 is not completely skipped but replaced by a learned vector, to inform the model that the chain is missing.

Because of the incompleteness of the data we want to learn as much as possible from every piece of data available. The decoder is in itself a language model, so it is able to learn without or with little conditioning. In a standard encoder-decoder Transformer learning the encoder is only trained through the decoder. We want to avoid this so that the encoder learning will not be entirely dependent on another piece of data to predict. Luckily, encoders are also trainable alone (without a decoder) by doing Masked Language Modeling (MLM). During MLM training we pick 15% of the amino acid positions, we’ll call this set of amino acids *ℳ*. From these ones 80 % are replaced by a mask token, and 20 % are replaced by a random amino acids. This deteriorated sequence is fed to the encoder. We learn a linear classifier on top to predict the original amino acids in *ℳ*. The logits output by this classifier are then passed by a softmax, defining for every position *i* a distribution over the amino acids *a*_*i*_: *P*_*cls*_(*a*_*i*_|*z*_*masked*_). The final MLM loss is simply the cross entropy:

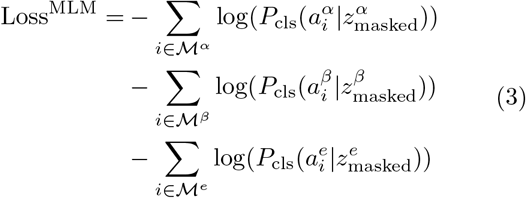

For example, it enables the epitope-encoder to still learn from the 600000 samples where we do not have TCRs.

In the end we combined the two losses, using a parameter *λ* that we always use equal to 0.5 in this paper, and sum over the sequence in the training set:

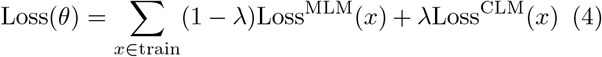

where *x* = (*x*_*α*_, *x*_*β*_, *x*_*e*_, MHC) is our raw datapoint and *θ* are the parameters of the model. Details on the training of a transformer can be found in appendix.

Code and weights for the model can be found at https://github.com/barthelemymp/TULIP-TCR/

### C. Mutual information as a proxy to the binding probability

The model presented before is autoregressive. The structure of the probabilities defined by the model is simply 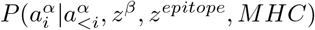(resp *β*, epitope) and a simple multiplication over the position gives us a conditional probability on the sequences (*p*(*e*|*α, β*, MHC), p(*α*|e, *β*, MHC), p(*β*|e, *α*, MHC). However, we should be more precise on what we want to evaluate. These conditional probabilities can be good for generating sequences, but here we first want to evaluate the probability of binding. We will show in this section how to approximate this quantity from the ones evaluated by our model. Let’s introduce the random binary variable of binding or not *b* such that *e T* becomes dependant conditionally on *b*. The first thing we need to observe is that our TULIP model is trained only on positive, i.e., binding examples. As a first simplification let’s look at the link between the binding posterior for a simple case of *e* being the epitope and *T* the alpha and beta chain of the TCR A simple bayesian approach will help us here.

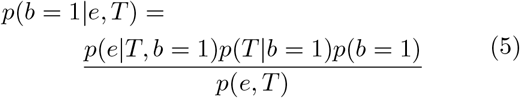

we can start to do some approximation here.

- All TCR sequenced in blood should have passed some positive thymic selection for epitope binding. This implies that *p*(*T* |*b* = 1) = *p*(*T*)
- *p*(*e, T*) = *p*(*e*)*p*(*T*) by construction as the dependence only appears when conditioning on b.

Leading to:

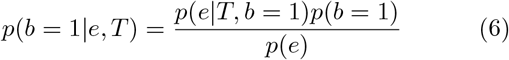

noticing the *p*(*b* = 1) are constants of the problem, we see that the binding posterior is proportional to the pointwise mutual information (PMI) between *T* and *e*:

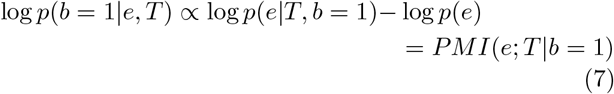

This quantity is the one we used to validate our models in the previous sections. Pushing further the derivation to include the role of the MHC, did not improve the results.

A similar computation can be done for the interaction between the epitope and MHC, by simply replacing T with MHC in the previous equation. This second term is used in Section. II D, where the experimental EC50 are the results of the simultaneous binding of the TCR with epitope and of the epitope with the MHC.

## V. DATA AND CODE AVAILABILITY

Code is available at barthelemymp/TULIP-TCR/. https://github.com/ The data used were collected from https://vdjdb.cdr3.net/, https://www.iedb.org/ and http://friedmanlab.weizmann.ac.il/McPAS-TCR/.

## ACKNOWLEDGMENTS

A.M.W. and T.M. were supported by grant COG 724208 from the European Research Council, and grant ANR-19-CE45-0018 “RESP-REP” from the Agence Nationale de la Recherche. This work was granted access to the HPC resources of IDRIS under the allocation 2022-AD011013872 made by GENCI.

## APPENDICES

### Appendix A Training details

In all the examples presented in this paper, we used the following architecture: embedding dim = 128, hidden size = 128, and each encoder and each decoder have 2 layers. The MHC embedding was limited to the 50 most represented MHC. In all the examples presented in this paper, we used the following architecture: embedding dimension = 128, hidden size = 128, and 2 layers for each encoder and each decoder. The MHC embedding was limited to the 50 most represented MHC, and none in the training with unseen epitopes. For the repertoire mining experiment, we used 100 epochs in the zero shot setting. During the training we use the Adam optimizer [45] with a learning rate of 0.0001 for 100 epochs. During the finetuning process, we freeze the encoder and the embedding. We train using the same losses with adam optimizer. The finetuning is done for 40 epochs (but on much smaller dataset) keeping the loss function the same.

### Appendix B: Comparison to other methods

For comparing with DLpTCR we fine-tuned our model on HLA-A02*01 in the same way as for the experiments of section II.A. For ERGO, metrics are computed on the subset of peptide that were both left out by TULIP and ERGO.

**TABLE S1.**
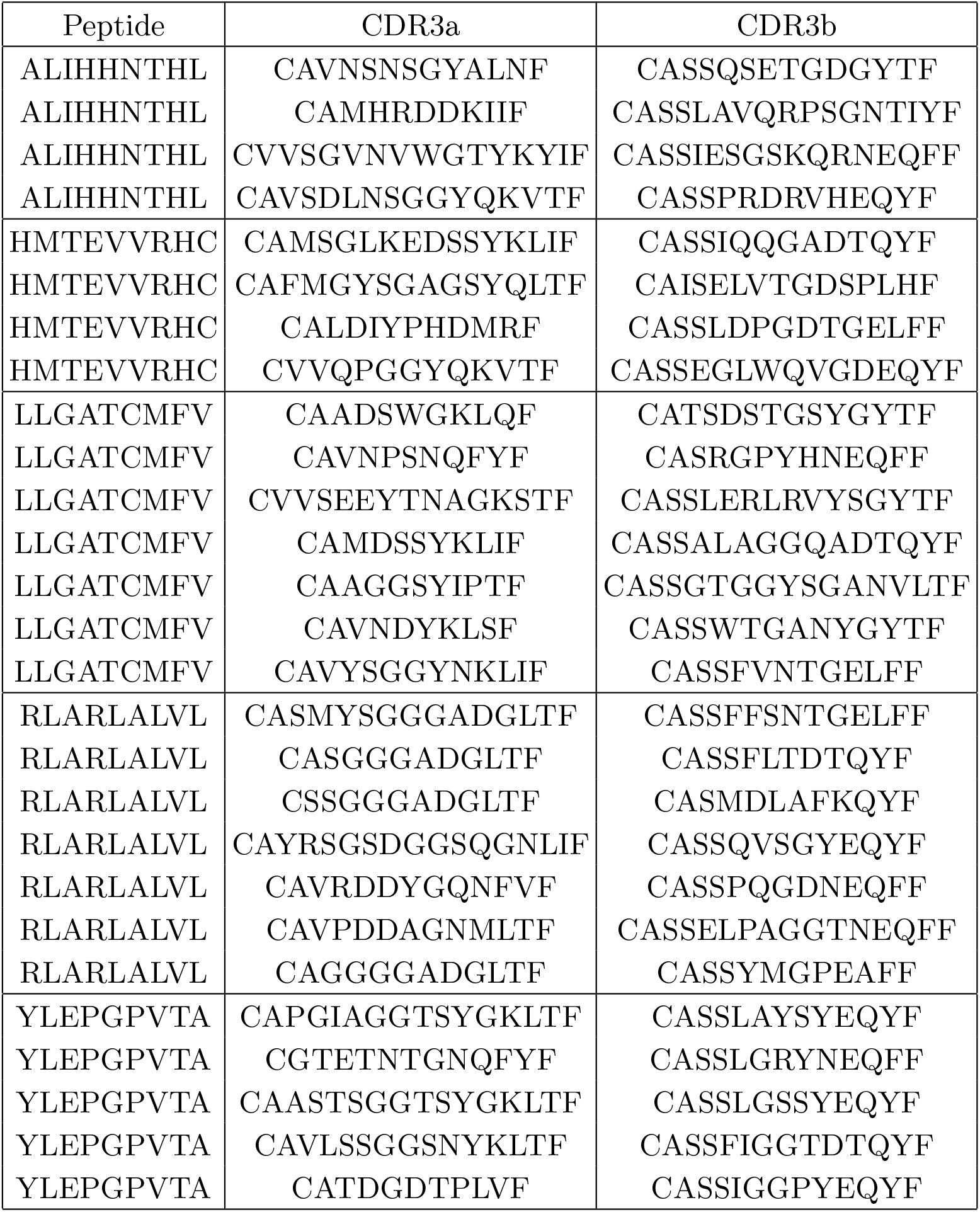
TCRs used in the repertoire mining tests.

**FIG. S1.**
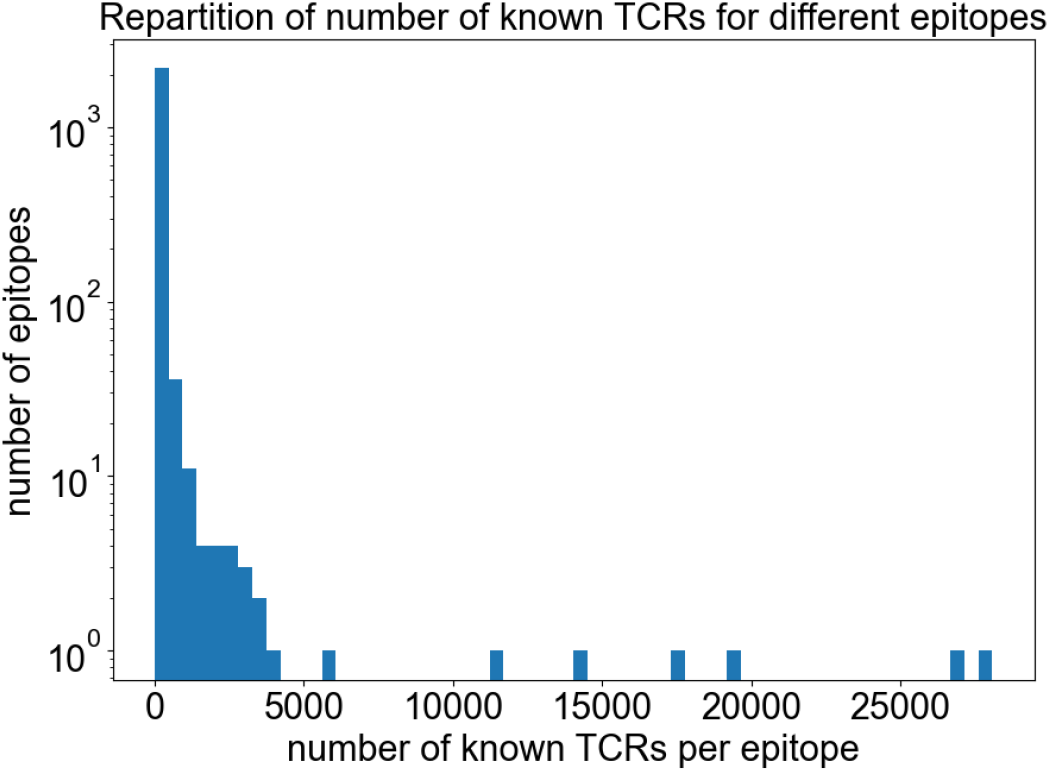
For each epitope in our database, we counted the number of known T-cell receptors (TCRs) binding to that epitope. The histogram shows a strong imbalance in the dataset, where a handful of epitopes harbor a substantial number of known TCRs, while the majority of epitopes have only a limited number of associated TCRs.

**FIG. S2.**
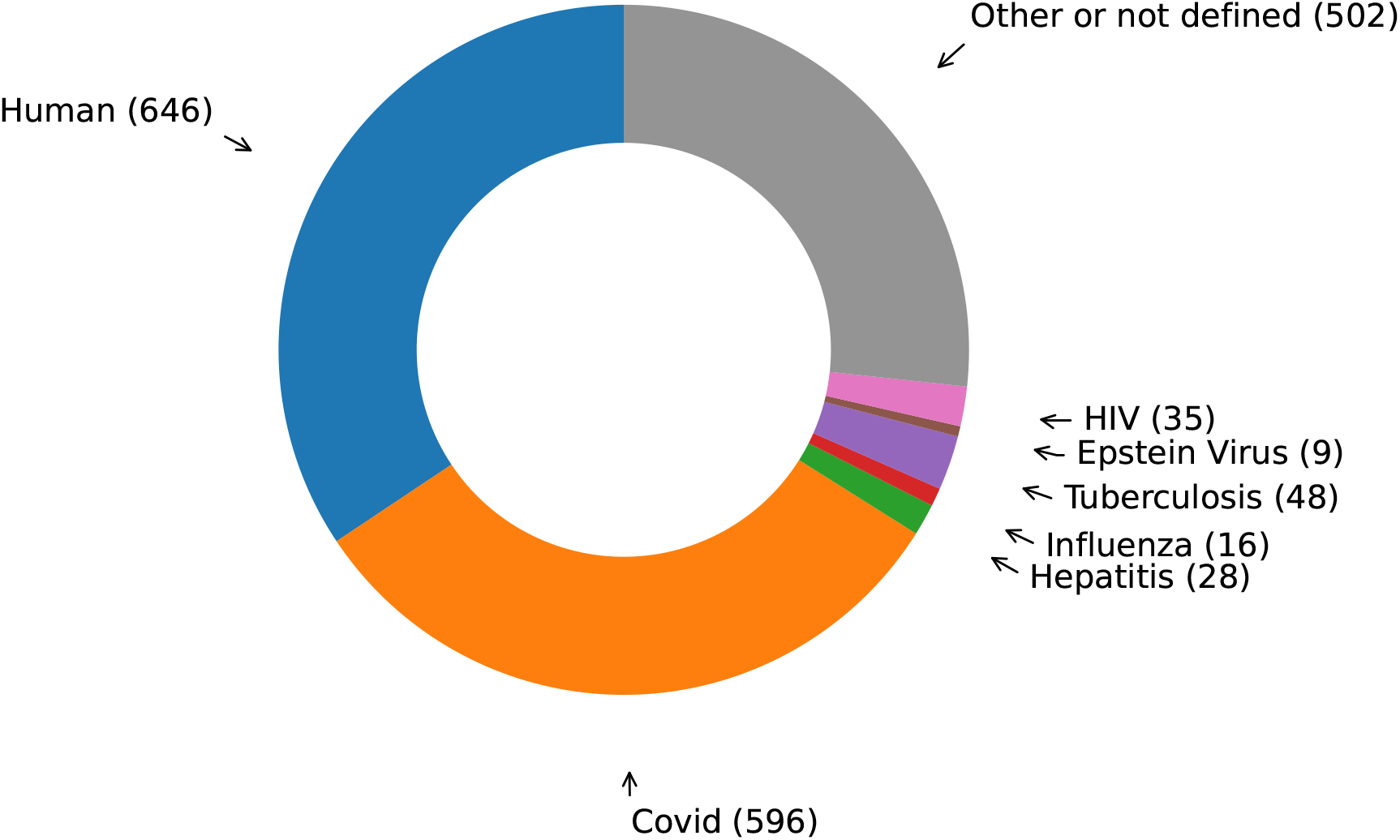
Distribution of peptides by organism of origin.

**FIG. S3.**
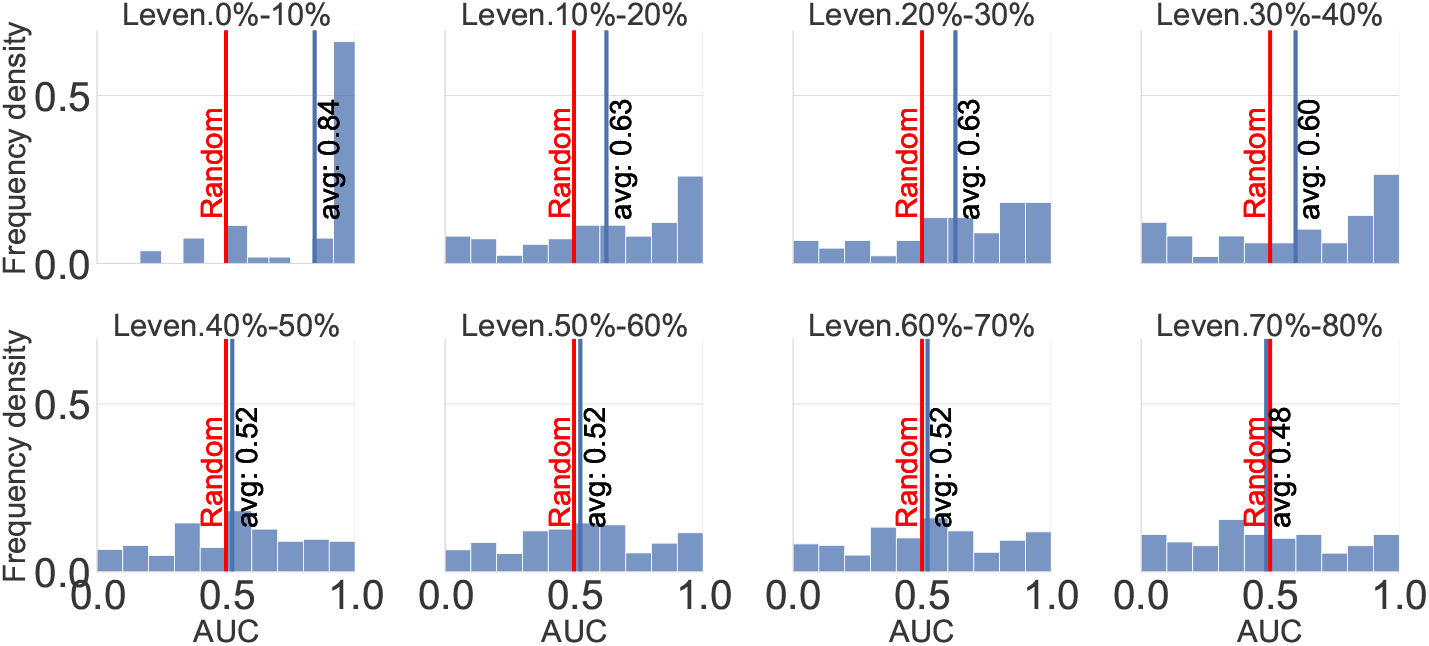
Distribution of the AUCs for the different distance groups. The plot complements Fig. 3A.

**FIG. S4.**
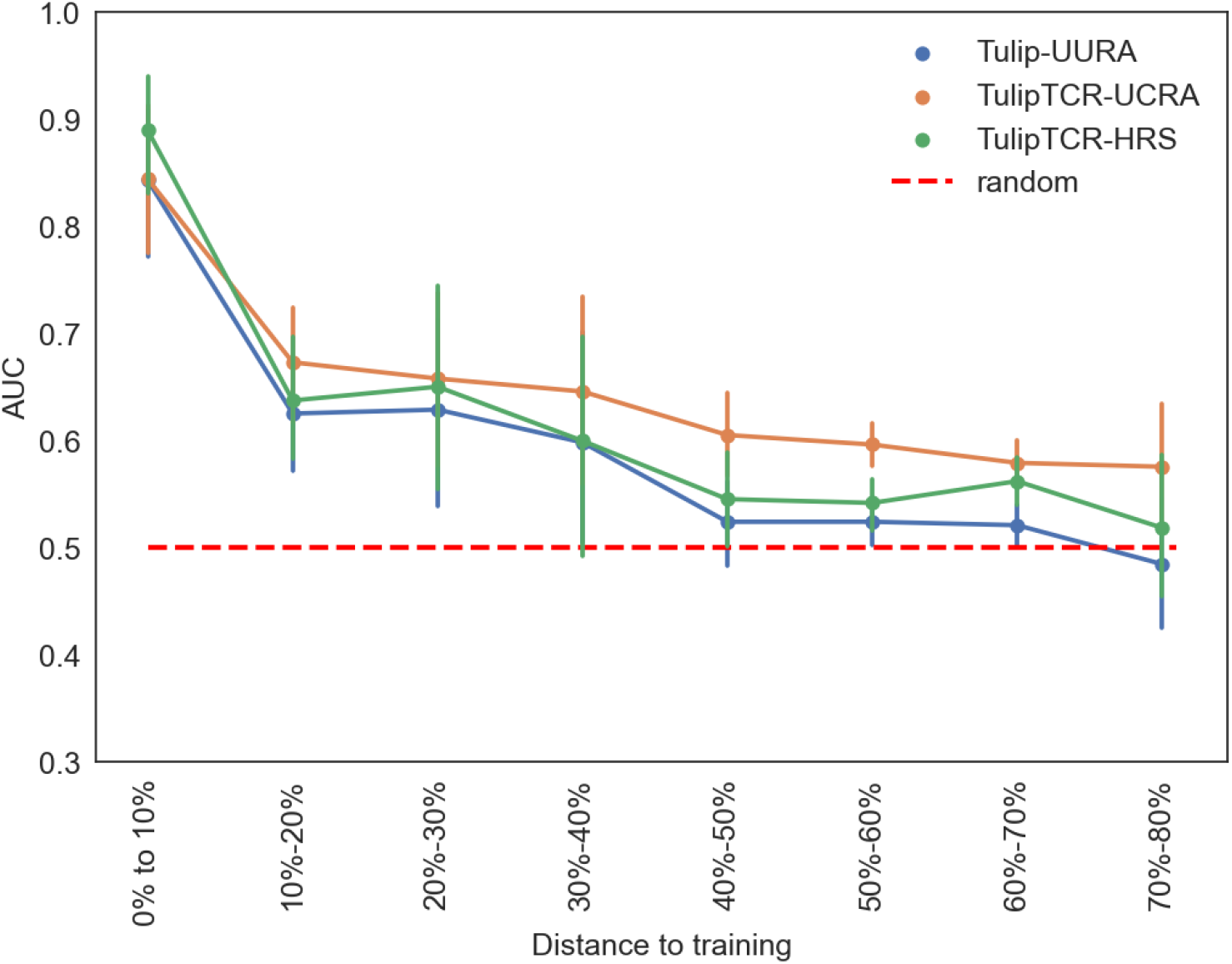
Mean AUC of the different distance groups. Equivalent to Fig. 3A, but with normalized distance (Hamming distance divided by length).

**FIG. S5.**
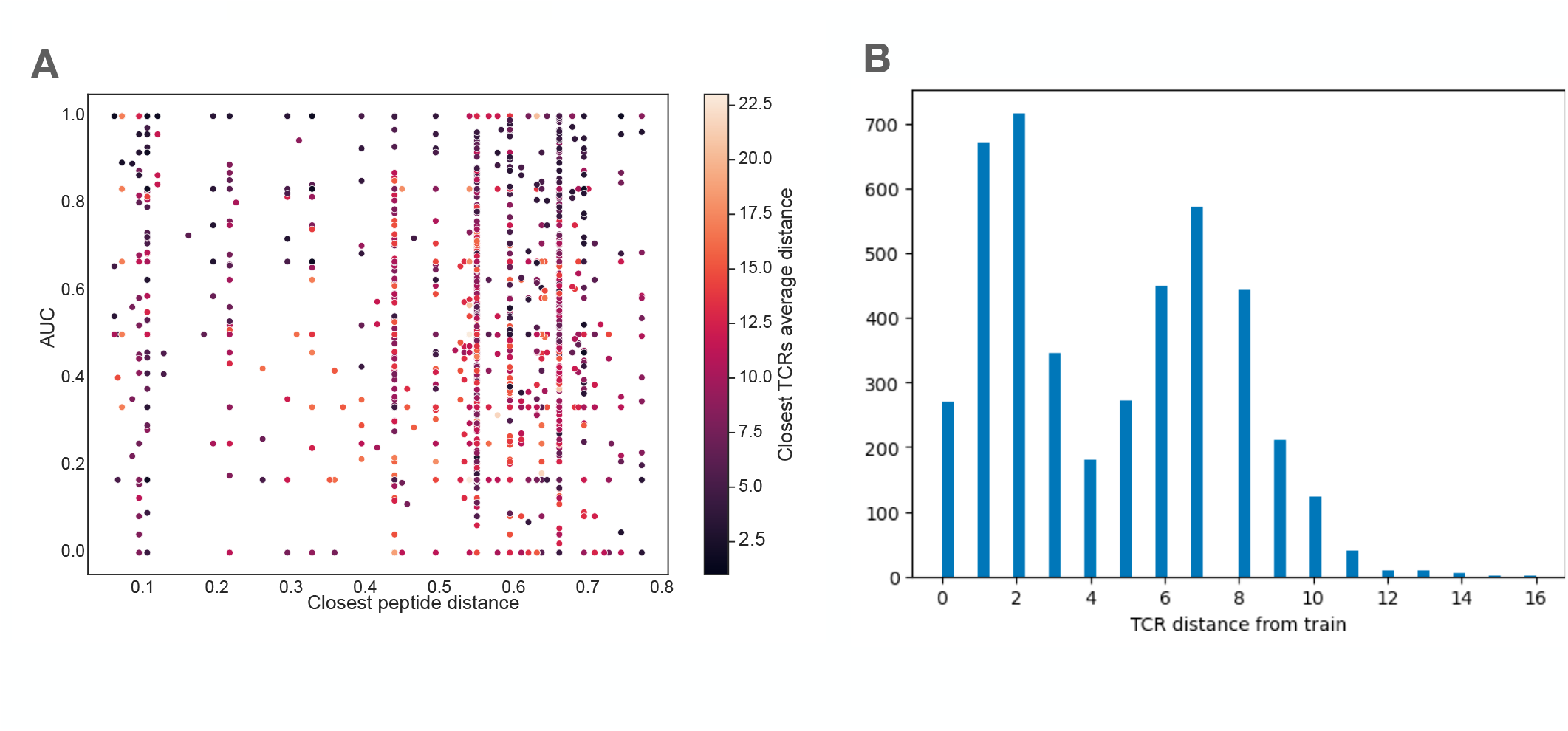
A. Scatter plot of AUC of individual peptides, versus the normalized distance (Hamming distance divided by CDR3 length) to closest peptide in the training set. The color indicates the average distance between a TCR associated to the peptide of interest, and the closest TCR associated to its closest peptides. B. Distribution of Hamming distances between a sequence from the test set and its closest TCR in the train set.

**FIG. S6.**
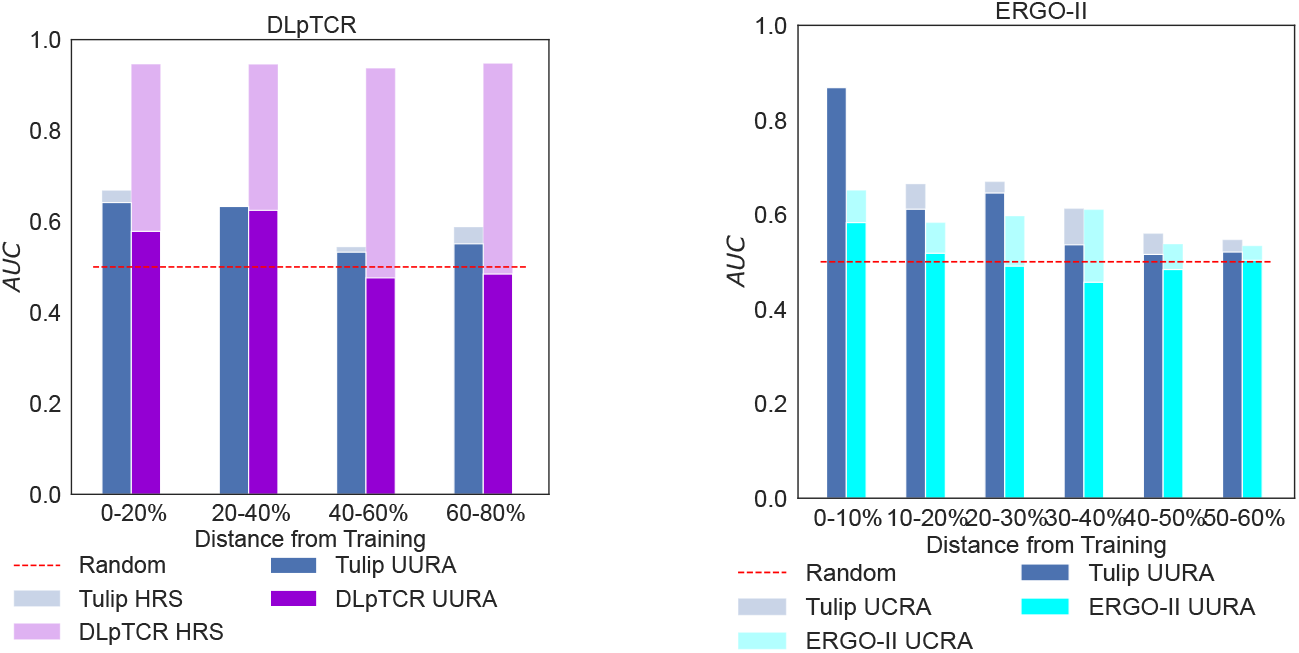
Performance of DLpTCR and ERGO2 on unseen peptides (complement to Fig. 3C).

**FIG. S7.**
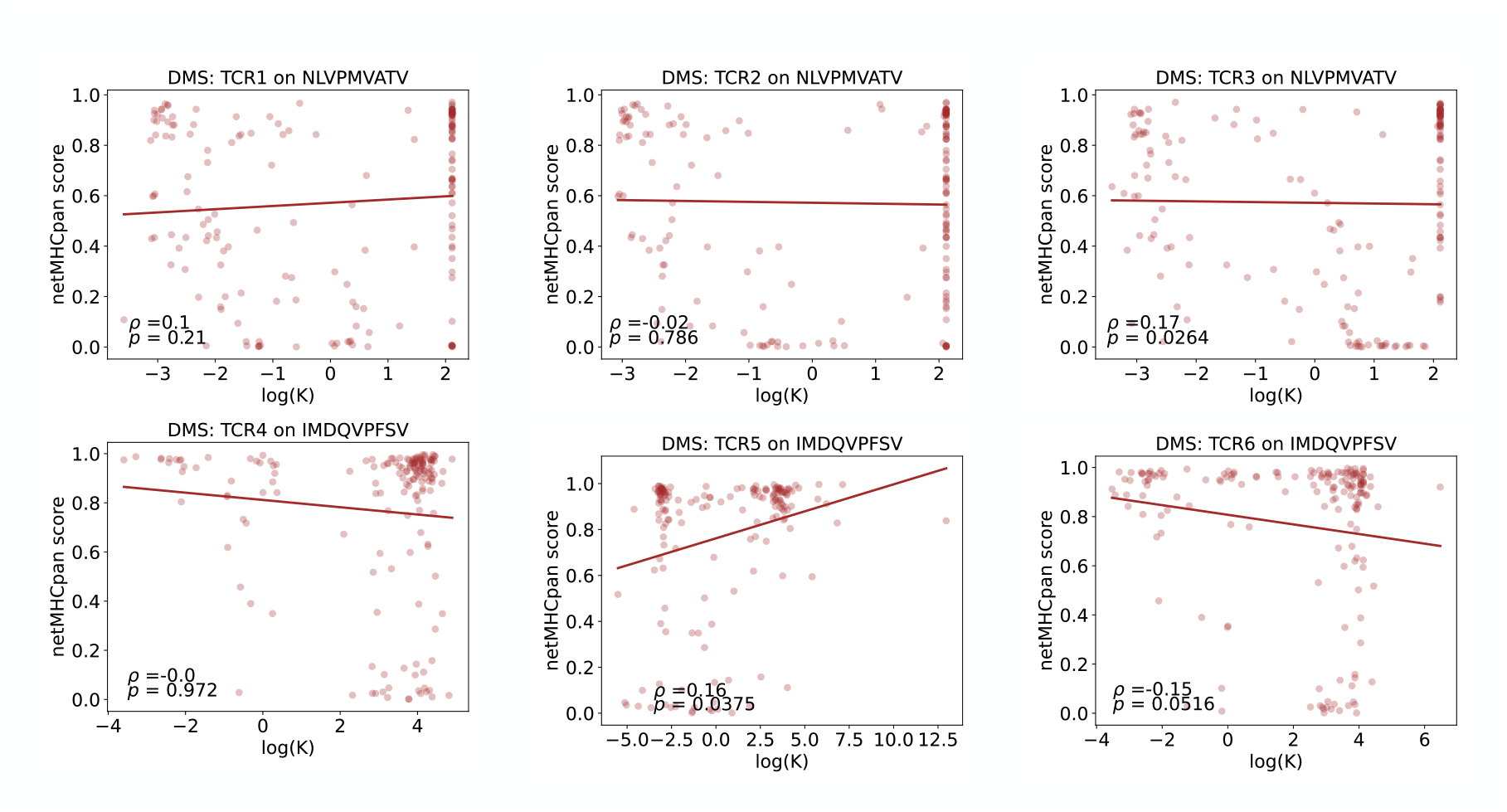
Effect of single epitope mutations on the NetMHCPan [36] score (log *P*) predicts TCR binding (dissociation constant *K*, in *μ*g.ml^*−*1^) measured by deep mutational scan experiments [35]. The reported *ρ* and p-values correspond to Spearman correlations.

## References

[1] O. Feinerman, J. Veiga, J. R. Dorfman, R. N. Germain, and G. Altan-Bonnet, Variability and robustness in t cell activation from regulated heterogeneity in protein levels, Science 321, 1081 (2008).

[2] P. François, G. Voisinne, E. D. Siggia, G. Altan-Bonnet, and M. Vergassola, Phenotypic model for early t-cell activation displaying sensitivity, specificity, and antagonism, Proceedings of the National Academy of Sciences 110, E888 (2013).

[3] M. Poorebrahim, N. Mohammadkhani, R. Mahmoudi, M. Gholizadeh, E. Fakhr, and A. Cid-Arregui, Tcr-like cars and tcr-cars targeting neoepitopes: An emerging potential, Cancer gene therapy 28, 581 (2021).

[4] L. A. Rojas, Z. Sethna, K. C. Soares, C. Olcese, N. Pang, E. Patterson, J. Lihm, N. Ceglia, P. Guasp, A. Chu, et al., Personalized rna neoantigen vaccines stimulate t cells in pancreatic cancer, Nature , 1 (2023).

[5] P. Bradley, Structure-based prediction of t cell receptor: peptide-mhc interactions, eLife 12, e82813 (2023).

[6] D. S. Shcherbinin, V. K. Karnaukhov, I. V. Zvyagin, D. M. Chudakov,, and M. Shugay, Large-scale template-based structural modeling of t-cell receptors with known antigen specificity reveals complementarity features, bioRxiv , 2023 (2023).

[7] P. Dash, A. J. Fiore-Gartland, T. Hertz, G. C. Wang, S. Sharma, A. Souquette, J. C. Crawford, E. B. Clemens, T. H. Nguyen, K. Kedzierska, et al., Quantifiable predictive features define epitope-specific t cell receptor repertoires, Nature 547, 89 (2017).

[8] J. Glanville, H. Huang, A. Nau, O. Hatton, L. E. Wagar, F. Rubelt, X. Ji, A. Han, S. M. Krams, C. Pettus, et al., Identifying specificity groups in the t cell receptor repertoire, Nature 547, 94 (2017).

[9] D. V. Bagaev, R. M. Vroomans, J. Samir, U. Stervbo, C. Rius, G. Dolton, A. Greenshields-Watson, M. Attaf, E. S. Egorov, I. V. Zvyagin, et al., Vdjdb in 2019: database extension, new analysis infrastructure and a t-cell receptor motif compendium, Nucleic Acids Research 48, D1057 (2020).

[10] R. Vita, S. Mahajan, J. A. Overton, S. K. Dhanda, S. Martini, J. R. Cantrell, D. K. Wheeler, A. Sette,, and B. Peters, The immune epitope database (iedb): 2018 update, Nucleic acids research 47, D339 (2019).

[11] N. Tickotsky, T. Sagiv, J. Prilusky, E. Shifrut, and N. Friedman, Mcpas-tcr: a manually curated catalogue of pathology-associated t cell receptor sequences, Bioinformatics 33, 2924 (2017).

[12] T. Mora and A. M. Walczak, Quantifying lymphocyte receptor diversity, in Systems Immunology (CRC Press, 2018) pp. 183–198.

[13] P. Meysman, J. Barton, B. Bravi, L. Cohen-Lavi, V. Karnaukhov, E. Lilleskov, A. Montemurro, M. Nielsen, T. Mora, P. Pereira, et al., Benchmarking solutions to the t-cell receptor epitope prediction problem: Immrep22 workshop report, ImmunoInformatics 9, 100024 (2023).

[14] A. Montemurro, V. Schuster, H. R. Povlsen, A. K. Bentzen, V. Jurtz, W. D. Chronister, A. Crinklaw, S. R. Hadrup, O. Winther, B. Peters, et al., Nettcr-2.0 enables accurate prediction of tcr-peptide binding by using paired tcrα and β sequence data, Communications biology 4, 1060 (2021).

[15] Y. Tong, J. Wang, T. Zheng, X. Zhang, X. Xiao, X. Zhu, X. Lai, and X. Liu, Sete: Sequence-based ensemble learning approach for tcr epitope binding prediction, Computational Biology and Chemistry 87, 107281 (2020).

[16] S. Gielis, P. Moris, N. De Neuter, W. Bittremieux, B. Ogunjimi, K. Laukens, and P. Meysman, Tcrex: a webtool for the prediction of t-cell receptor sequence epitope specificity, BioRxiv 373472 (2018).

[17] E. Jokinen, J. Huuhtanen, S. Mustjoki, M. Heinonen, and H. Lähdesmäki, Predicting recognition between t cell receptors and epitopes with tcrgp, PLoS computational biology 17, e1008814 (2021).

[18] P. Dash, A. J. Fiore-Gartland, T. Hertz, G. C. Wang, S. Sharma, A. Souquette, J. C. Crawford, E. B. Clemens, T. H. Nguyen, K. Kedzierska, et al., Quantifiable predictive features define epitope-specific t cell receptor repertoires, Nature 547, 89 (2017).

[19] A. Weber, J. Born, and M. Rodriguez Martínez, Titan: T-cell receptor specificity prediction with bimodal attention networks, Bioinformatics 37, i237 (2021).

[20] Y. Gao, Y. Gao, Y. Fan, C. Zhu, Z. Wei, C. Zhou, G. Chuai, Q. Chen, H. Zhang, and Q. Liu, Pan-peptide meta learning for t-cell receptor–antigen binding recognition, Nature Machine Intelligence , 1 (2023).

[21] I. Springer, N. Tickotsky, and Y. Louzoun, Contribution of t cell receptor alpha and beta cdr3, mhc typing, v and j genes to peptide binding prediction, Frontiers in immunology 12, 664514 (2021).

[22] B. P. Kwee, M. Messemaker, E. Marcus, G. Oliveira, W. Scheper, C. Wu, J. Teuwen, and T. Schumacher, Stapler: Efficient learning of tcr-peptide specificity prediction from full-length tcr-peptide data, bioRxiv , 2023 (2023).

[23] Z. Xu, M. Luo, W. Lin, G. Xue, P. Wang, X. Jin, C. Xu, W. Zhou, Y. Cai, W. Yang, et al., Dlptcr: an ensemble deep learning framework for predicting immunogenic peptide recognized by t cell receptor, Briefings in Bioinformatics 22, bbab335 (2021).

[24] E. Lanzarotti, P. Marcatili, and M. Nielsen, T-cell receptor cognate target prediction based on paired α and β chain sequence and structural cdr loop similarities, Frontiers in immunology 10, 2080 (2019).

[25] J. J. Moon, H. H. Chu, M. Pepper, S. J. McSorley, S. C. Jameson, R. M. Kedl, and M. K. Jenkins, Naive cd4+ t cell frequency varies for different epitopes and predicts repertoire diversity and response magnitude, Immunity 27, 203 (2007).

[26] M. Yousef, S. Jung, L. C. Showe, and M. K. Showe, Learning from positive examples when the negative class is undetermined-microrna gene identification, Algorithms for molecular biology 3, 1 (2008).

[27] P. Perera, P. Oza, and V. M. Patel, One-class classification: A survey, arXiv preprint arXiv:2101.03064 (2021).

[28] T. Brown, B. Mann, N. Ryder, M. Subbiah, J. D. Kaplan, P. Dhariwal, A. Neelakantan, P. Shyam, G. Sastry, A. Askell, et al., Language models are few-shot learners, Advances in neural information processing systems 33, 1877 (2020).

[29] A. Radford, K. Narasimhan, T. Salimans, I. Sutskever, et al., Improving language understanding by generative pre-training, OpenAI (2018).

[30] A. Vaswani, N. Shazeer, N. Parmar, J. Uszkoreit, L. Jones, A. N. Gomez, L. Kaiser, and I. Polosukhin, Attention is all you need, Advances in neural information processing systems 30 (2017).

[31] B. Meynard-Piganeau, C. Fabbri, M. Weigt, A. Pagnani, and C. Feinauer, Generating interacting protein sequences using domain-to-domain translation, bioRxiv , 2022 (2022).

[32] Z. Wu, K. K. Yang, M. J. Liszka, A. Lee, A. Batzilla, D. Wernick, D. P. Weiner, and F. H. Arnold, Signal peptides generated by attention-based neural networks, ACS Synthetic Biology 9, 2154 (2020).

[33] K. C. Garcia and E. J. Adams, How the t cell receptor sees antigen—a structural view, Cell 122, 333 (2005).

[34] H. Tanno, T. M. Gould, J. R. McDaniel, W. Cao, Y. Tanno, R. E. Durrett, D. Park, S. J. Cate, W. H. Hildebrand, C. L. Dekker, et al., Determinants governing t cell receptor α/β-chain pairing in repertoire formation of identical twins, Proceedings of the National Academy of Sciences 117, 532 (2020).

[35] M. Luksza, Z. M. Sethna, L. A. Rojas, J. Lihm, B. Bravi, Y. Elhanati, K. Soares, M. Amisaki, A. Dobrin, D. Hoyos, et al., Neoantigen quality predicts immunoediting in survivors of pancreatic cancer, Nature 606, 389 (2022).

[36] B. Reynisson, B. Alvarez, S. Paul, B. Peters, and M. Nielsen, Netmhcpan-4.1 and netmhciipan-4.0: improved predictions of mhc antigen presentation by concurrent motif deconvolution and integration of ms mhc eluted ligand data, Nucleic acids research 48, W449 (2020).

[37] E. Kondo, B. Maecker, M. R. Weihrauch, C. Wickenhauser, W. Zeng, L. M. Nadler, J. L. Schultze, and M. S. von Bergwelt-Baildon, Cyclin d1–specific cytotoxic t lymphocytes are present in the repertoire of cancer patients: Implications for cancer immunotherapy, Clinical Cancer Research 14, 6574 (2008).

[38] P. Malekzadeh, A. Pasetto, P. F. Robbins, M. R. Parkhurst, B. C. Paria, L. Jia, J. J. Gartner, V. Hill, Z. Yu, N. P. Restifo, et al., Neoantigen screening identifies broad tp53 mutant immunogenicity in patients with epithelial cancers, The Journal of clinical investigation 129 (2021).

[39] D. Wu, R. Gowathaman, B. G. Pierce, and R. A. Mariuzza, T cell receptors employ diverse strategies to target a p53 cancer neoantigen, Journal of Biological Chemistry 298 (2022).

[40] A. Scardino, D.-A. Gross, P. Alves, J. L. Schultze, S. Graff-Dubois, O. Faure, S. Tourdot, S. Chouaib, L. M. Nadler, F. A. Lemonnier, et al., Her-2/neu and htert cryptic epitopes as novel targets for broad spectrum tumor immunotherapy, The Journal of Immunology 168, 5900 (2002).

[41] S. S. Tykodi, S. Satoh, J. D. Deming, J. Chou, R. Harrop, and E. H. Warren, Cd8+ t cell clones specific for the 5t4 antigen target renal cell carcinoma tumor-initiating cells in a murine xenograft model, Journal of immunotherapy (Hagerstown, Md.: 1997) 35, 523 (2012).

[42] A. L. Cox, J. Skipper, Y. Chen, R. A. Henderson, T. L. Darrow, J. Shabanowitz, V. H. Engelhard, D. F. Hunt, and C. L. Slingluff Jr, Identification of a peptide recognized by five melanoma-specific human cytotoxic t cell lines, Science 264, 716 (1994).

[43] A. A. Minervina, E. A. Komech, A. Titov, M. Bensouda Koraichi, E. Rosati, I. Z. Mamedov, A. Franke, G. A. Efimov, D. M. Chudakov, T. Mora, et al., Longitudinal high-throughput tcr repertoire profiling reveals the dynamics of t-cell memory formation after mild covid-19 infection, Elife 10, e63502 (2021).

[44] A. S. Shomuradova, M. S. Vagida, S. A. Sheetikov, K. V. Zornikova, D. Kiryukhin, A. Titov, I. O. Peshkova, A. Khmelevskaya, D. V. Dianov, M. Malasheva, et al., Sars-cov-2 epitopes are recognized by a public and diverse repertoire of human t cell receptors, Immunity 53, 1245 (2020).

[45] D. P. Kingma and J. Ba, Adam: A method for stochastic optimization, arXiv preprint arXiv:1412.6980 (2014).

